# HARVEST: Unlocking the Dark Bioactivity Data of Pharmaceutical Patents via Agentic AI

**DOI:** 10.64898/2026.03.15.711910

**Authors:** Viktoriia Shepard, Aibulat Musin, Kristina Chebykina, Natalia A. Zeninskaya, Lukia Mistryukova, Konstantin Avchaciov, Peter O. Fedichev

**Affiliations:** GERO PTE. LTD., 133 Cecil Street 14-01 Keck Seng Tower, Singapore 069535

## Abstract

Pharmaceutical patents contain vast Structure–Activity Relationship tables documenting protein– ligand binding data. While technically public, this information remains computationally inaccessible and effectively dark, trapped in bulky documents that no existing database has systematically captured. We present HARVEST, a multi-agent large language model pipeline that autonomously extracts structured bioactivity records from USPTO patent archives at $0.11 per document. Applied to 164,877 patents, HARVEST produced 3.15 million activity records, recovering 326,342 unique scaffolds and 967 protein targets absent from BindingDB. This pipeline completed in under a week a task that would otherwise require over 55 years of continuous expert labor. Automated extraction achieves 80% agreement with human curated corpus of US patents from BindingDB, a conservative lower bound given identified errors within the reference data. We further introduce H-Bench, a structurally guaranteed held-out benchmark built from this recovered data. Evaluation of the leading open-source model Boltz-2 on H-Bench reveals a two-dimensional generalization gap: performance degrades both on novel scaffolds and on uncharacterized protein targets, exposing fundamental limitations of models trained on existing public repositories.

## I. INTRODUCTION

Pharmaceutical patents represent one of the largest repositories of experimental protein–ligand interaction (PLI) data ever assembled. Thousands of Structure– Activity Relationship (SAR) tables, each documenting binding affinities across hundreds of compounds, are filed annually with patent offices worldwide and often appear years before or independently of peer-reviewed literature [1–3]. Despite billions of dollars in R&D investment, this knowledge remains effectively “dark”: technically public, yet computationally inaccessible, trapped in unstructured archives that no existing database has systematically captured.

This data gap matters acutely. Recent breakthroughs in *de novo* protein design [4–6] and structure prediction [7–9] have transformed what AI can do in drug discovery, but these models face a generalization crisis. Even the best architectures struggle to predict activity in new chemical or biological spaces when trained on sparse data [10–12]. Closing this gap requires two things simultaneously: massive, diverse training sets and genuinely held benchmarks to demonstrate robust model generalization [13–15]. Pharmaceutical patents could provide both provided their contents were easily accessible.

PLI data is the ground truth for both training and benchmarking [16, 17]. The leading public repository, BindingDB, relies on the slow manual curation of the literature [18] and covers only a fraction of the available patent data. Automating patent extraction has historically failed due to the specific challenges of patent language and their multimodal nature [19, 20]: information is fragmented across unstructured text, complex tables, and chemical diagrams, and high-fidelity extraction demands reconstructing the complete link between a specific compound, the assay performed, and the resulting activity against a protein target [20]. Existing pipelines like SureChEMBL index chemical structures at scale but lack systematic extraction of quantitative binding values or mapping to biological targets [21, 22]. The result is a self-reinforcing bottleneck: models are evaluated on the same datasets they were trained on, making it impossible to distinguish genuine generalization from memorization.

Agentic AI systems break this bottleneck. Decomposing complex extraction into specialized sequential agents reduces hallucination rates and maintains accurate compound–target associations across documents exceeding 500,000 tokens. With the rapid rise in LLM reasoning and falling inference costs [23, 24], hierarchies of specialized agents can now mimic expert human workflows at negligible marginal cost – making systematic patent mining economically feasible for the first time.

We present **HARVEST** (High-throughput Agent Retrieval of Values for Evaluated Small-molecules and Targets), an automated multi-agent pipeline for the extraction of SARs from USPTO bulk data. The system autonomously parses patent XML, resolves chemical aliases to canonical SMILES, and maps biological targets to UniProt identifiers. When applied to 164,877 patent archives, the pipeline produced 3.15 million activity records from 40,902 patents at a cost of only $0.11 per document – completing in under a week a task that would require over 55 years of continuous manual expert labor. This dataset substantially expands the known chemical-biological landscape, recovering 326,342 unique scaffolds and 967 protein targets entirely absent from BindingDB.

A central contribution of this work is **H-Bench**, an open benchmark derived from HARVEST comprising bioactivity data absent from all existing public repositories. H-Bench supports two distinct evaluation scenarios: scaffold-generalization on known targets, and target-generalization across proteins with no prior public bioactivity data. Our evaluation of the leading open-source structure-based model Boltz-2 [9] on H-Bench reveals a two-dimensional generalization gap: model performance degrades both when chemistry is novel and when protein targets lack prior bioactivity data – demonstrating that current models have not yet learned fully transferable binding physics. Together, HARVEST and H-Bench convert billions of dollars of inaccessible R&D knowledge into open scientific infrastructure, directly addressing the data bottleneck that limits AI-driven therapeutic discovery.

## II. RESULTS

### A. HARVEST Substantially Expands Public Bioactivity Space

The HARVEST pipeline (Section IX) was applied to 164,877 USPTO patent archives pre-selected for the presence of chemical structures and bioactivity mentions. Processing 50 documents in parallel, the system produced a final dataset of 3.15 million activity records extracted from 40,902 patents (25% of the input corpus), averaging 82 records per document containing extractable data. The remaining patents either lacked extractable bioactivity data or fell outside the pipeline’s current parsing capabilities (see Section IV). The automated pipeline achieves a consistently higher document throughput than manual curation across the full 25-year publication window examined (Fig. 1a). Although BindingDB reports a higher average number of activity records per patent (Fig. 1b), this is a result of human curators prioritizing the most data-rich documents. HARVEST achieves comparable yield on shared patents, confirming equivalent extraction depth. Because the marginal cost of processing is negligible, HARVEST captures data from thousands of documents that would not justify the cost of manual labor.

**FIG. 1:**
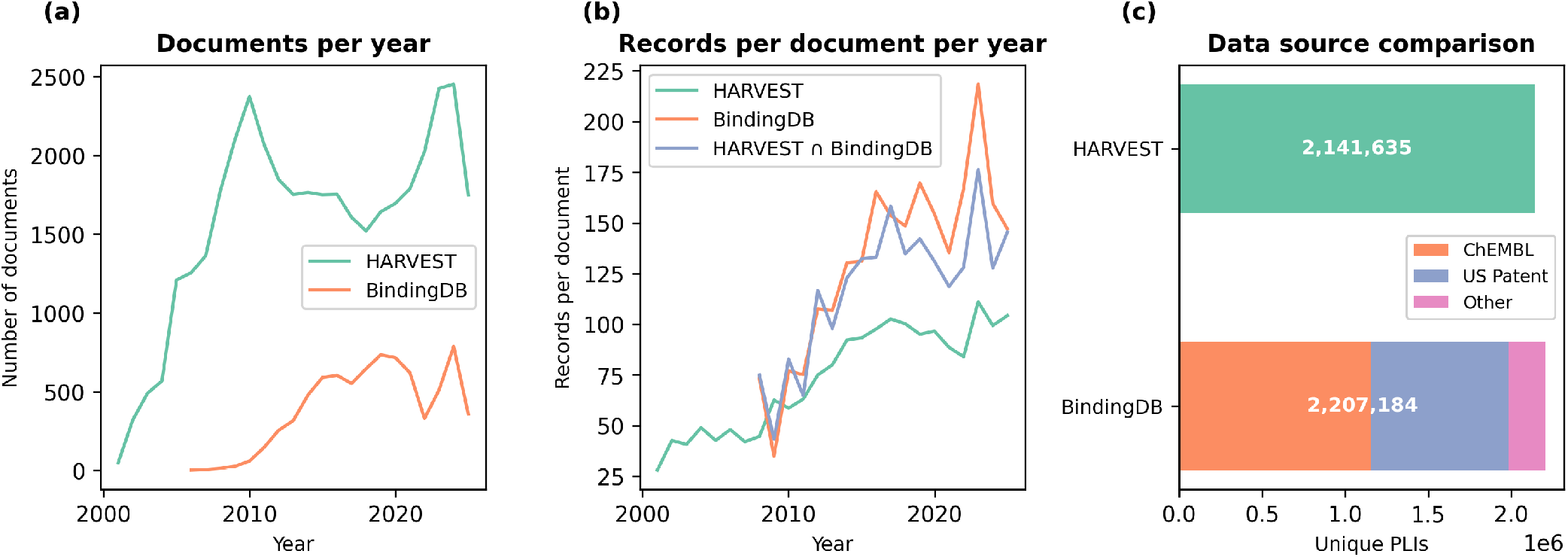
(a) Number of patents included per year from USPTO for HARVEST (green) and BindingDB (orange), illustrating the consistently higher document coverage achieved by automated extraction across the full 25-year window. (b) Mean number of activity records per document for HARVEST (green), BindingDB (orange), and their intersection (blue). The convergence of HARVEST and BindingDB on shared patents confirms equivalent extraction depth; the lower overall HARVEST average reflects broader document selection criteria that include patents with fewer activity records. (c) Comparison of unique protein–ligand interactions (PLIs) between HARVEST and BindingDB. The BindingDB bar is decomposed by data source: ChEMBL-derived entries (orange), US Patent extractions (blue), and remaining sources (pink). HARVEST yields 2.14M unique PLIs from patent text alone, nearly matching the 2.21M across all BindingDB sources combined.

Overall, HARVEST and BindingDB contain a comparable total volume of protein–ligand interactions. After aggregating multiple measurements per compound– target pair and applying inclusion filters (Section IX D), HARVEST yields 2.14M unique PLIs, comparable to BindingDB’s 2.21M across all sources (Fig. 1c). However, these totals reflect fundamentally different source compositions: BindingDB aggregates records from patents, journal articles, and ChEMBL, whereas HARVEST draws exclusively from patent text. Restricting the comparison to patent-derived records, HARVEST contributes nearly three times as many PLIs as BindingDB’s patent subset, confirming substantially deeper coverage of the patent corpus.

Across the combined chemical-biological landscape of 8,667 protein targets, only 35.1% are shared between the two databases (Fig. 2a). A further 967 targets (12.7%) are covered exclusively by HARVEST, while the majority of BindingDB-only targets (53.8%) reflect literature and ChEMBL sources outside the patent corpus. Critically, novelty extends well beyond HARVEST-exclusive proteins: for the 3,042 shared targets, 66.2% of HARVEST PLIs (1,369,936 interactions) and 46.0% of scaffolds (304,516 clusters) are absent from BindingDB (Fig. 2b–c). This indicates that substantial chemical diversity remains trapped in patent archives even for the most extensively studied drug targets.

**FIG. 2:**
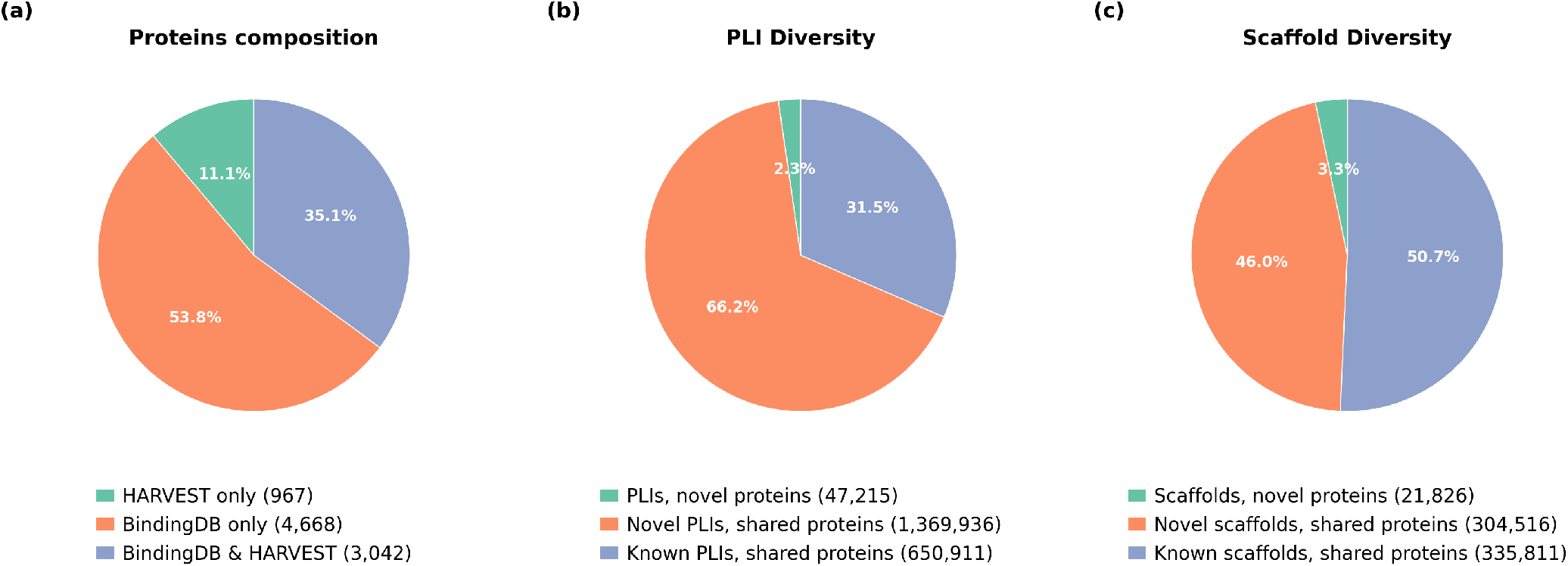
Comparison of protein and chemical space coverage between HARVEST and BindingDB. (a) Protein target composition across both databases (n = 8,667 total): 35.1% of targets are shared, while 11.1% are covered exclusively by HARVEST and 53.8% exclusively by BindingDB. (b) Protein–ligand interaction (PLI) diversity within HARVEST for the 3,042 shared protein targets: 66.2% of PLIs are novel relative to BindingDB, with an additional 2.3% associated with proteins unique to HARVEST. (c) Scaffold diversity within HARVEST for shared targets: 46.0% of scaffolds are absent from BindingDB, with a further 3.3% belonging to HARVEST-exclusive proteins. Orange indicates novel content unique to HARVEST; blue indicates overlap with BindingDB; green indicates targets absent from BindingDB entirely.

The target distribution reflects established drug discovery priorities: enzymes and kinases predominate due to their well-defined binding pockets [25], while transcription factors are less represented due to the difficulty of targeting protein-protein interfaces with small molecules (see Table S1).

To assess the utility of this expanded chemical space for structure-activity relationship (SAR) analysis, we quantified the density of activity cliffs. These are defined as pairs of structurally similar compounds (Tanimoto similarity *≥*0.7) that exhibit a substantial difference in biological activity (ΔpActivity *≥* 1.5), where pActivity = − log_10_[M] [26]. Activity cliffs represent high-information data points where minor chemical modifications critically determine a molecule’s effect. Across the 3,042 shared targets with at least one activity cliff, HARVEST provides greater cliff density for 42% of proteins, while BindingDB leads for 54%; only 3% show equivalent coverage. This asymmetric complementarity– where each resource uniquely enriches SAR information for distinct protein subsets–argues strongly for merging both datasets to maximize the structural discontinuities available for lead optimization and machine learning model training.

### B. HARVEST Extracts Data Comparable in Quality to Manual Curation

To validate the fidelity of automated extraction, we benchmarked HARVEST against the manually curated BindingDB (BDB) dataset [18]. Since HARVEST operates exclusively on US patents, all comparisons use the patent-derived subset of BDB unless otherwise noted.

Density distributions of binding affinity, molecular weight, and synthetic accessibility show close agreement between HARVEST and BDB across the full range of values (Figs. 3a–3c consistent with previously reported patent-derived compound profiles [18]. We next assessed record-level accuracy by pairwise comparison of activity values for PLIs present in both databases, matched by UniProt accession and InChIKey connectivity layer (first block) [27]. Across 319,954 matched PLIs from 5,668 shared patents, the distribution of activity residuals (ΔpActivity) is highly centered around zero, with 91.0% of PLIs showing near-identical values (Fig. 4a). This corresponds to a high quantitative correlation (Pearson *r* = 0.925, Spearman *ρ* = 0.937). Validation against 68,209 independent article-derived records confirms this consistency (*r* = 0.851, *ρ* = 0.875), with the broader residual distribution reflecting inherent experimental variability between separate data sources (Fig. 4b).

**FIG. 3:**
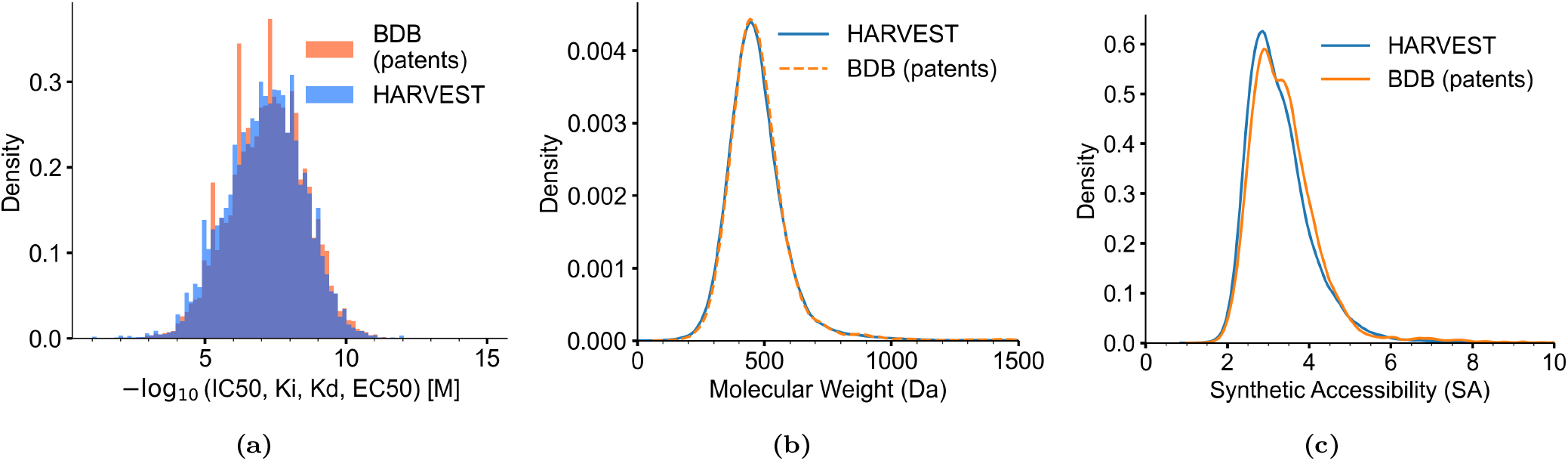
Physicochemical property distributions for HARVEST and BindingDB (US patents subset). (a) Binding affinity distributions (combined IC_50_, *K*_*i*_, *K*_*d*_, and EC_50_). Only exact numeric measurements (relation “=“) are included. (b) Molecular weight distributions, with both peaking near 450 Da. (c) Synthetic accessibility (SA) scores, both peaking at SA ≈ 3. The alignment across three descriptors confirms that HARVEST extracts a representative chemical space without systematic physicochemical bias relative to manual curation.

**FIG. 4:**
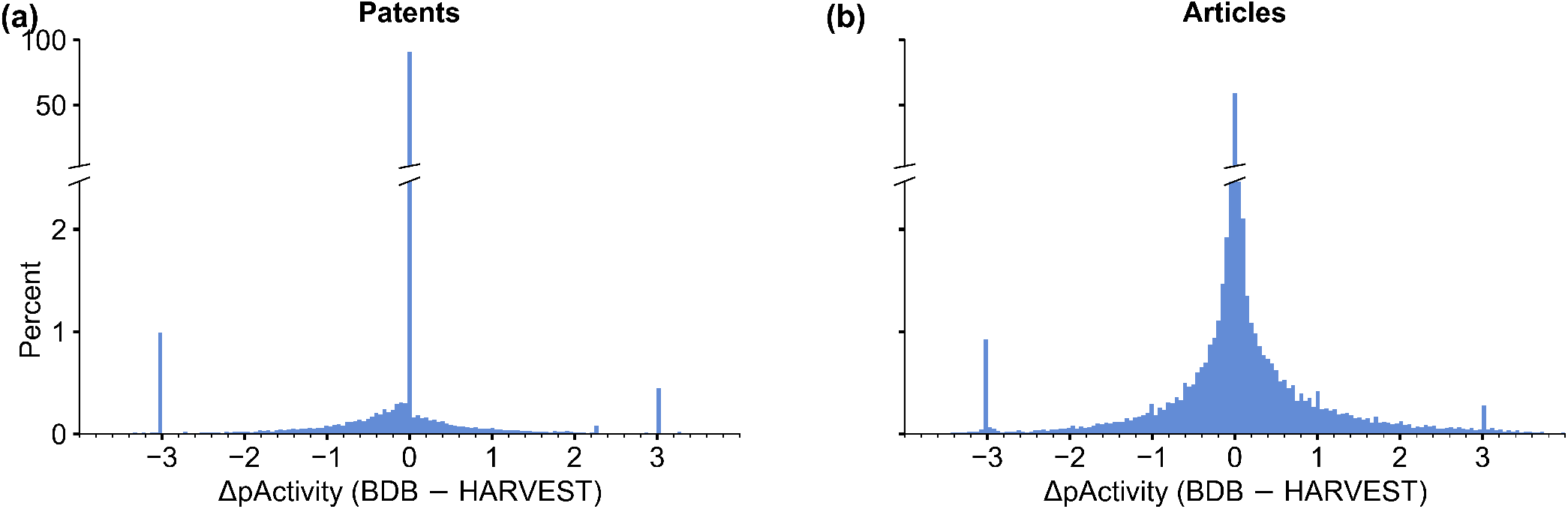
Distribution of activity residuals (ΔpActivity = BDB − HARVEST) for matched compound–target pairs. The y-axis is broken to show both the dominant central peak and the tail structure. (a) US patent-derived BindingDB records (*n* = 319,954): 91.0% of pairs fall in the central bin (ΔpActivity ≈ 0), corresponding to the high correlation (*r* = 0.925) observed between manual and automated curation. (b) Article-derived BindingDB records (*n* = 68,209): the broader distribution (59.3% central bin, *r* = 0.851) reflects genuine measurement variability across independent data sources rather than extraction error. In both panels, the distinct spikes at ΔpActivity = ±3 identify 1,000-fold unit conversion errors (nM/µM confusion), the most frequent artifact in both curation workflows. The sharp central distribution confirms that HARVEST achieves human-level extraction fidelity across hundreds of thousands of records.

The residual analysis reveals isolated spikes at ΔpActivity = ±3, the signature of 1,000-fold unit conversion errors (nM/µM confusion), the most common curation artifact reported by the BindingDB team [18]. These affect ∼1.4% of patent PLIs and ∼1.2% of article PLIs. To determine which database held the correct value, we manually verified the original patent text for the 20 patents contributing the most discrepant records, collectively covering 3,706 of the 5,499 affected PLIs (67%). At the record level, HARVEST held the correct value in 92% of verified PLIs, BindingDB in 5%, with 3% remaining ambiguous. This indicates that automated agents are substantially less prone to unit-conversion errors than manual curators.

The preceding comparison is restricted to records present in both databases. To assess record-level overlap, we cross-referenced each record against the other database within the same patent (Table I). Of 606,456 HARVEST records, 80.3% find an exact match in BindingDB; conversely, HARVEST recovers 70.5% of 690,869 BindingDB records. To understand the sources of discrepancy, we manually verified patents contributing the most records in each mismatch category (Tables S3–S5), collectively covering 12.1% of all mismatched records.

**TABLE I:** Cross-validation on 5,668 shared patents. Each row indicates how a record from one database was matched in the other on the same patent. *Full match*: both compound and target agree. *Target mismatch*: same compound found but assigned to a different protein. *Compound mismatch*: same target found but linked to a different compound. *No overlap*: neither compound nor target found on that patent.

|  | **HARVEST**<br>(*n*=606,456)<br>in BDB | **BDB**<br>(*n*=690,869)<br>in HARVEST |
| --- | --- | --- |
| Full match | 486,950 (80.3%) | 486,950 (70.5%) |
| Target mismatch | 61,823 (10.2%) | 53,784 (7.8%) |
| Compound mismatch | 47,685 (7.9%) | 124,061 (18.0%) |
| No overlap | 9,998 (1.6%) | 26,074 (3.8%) |

#### Target mismatch

When both databases find the same compound but assign it to different proteins, HARVEST more often identified the correct target. In the remaining cases, HARVEST misclassified assay readout biomarkers as direct binding targets. Additional disagreements arose from differences in protein name resolution and from unclear species assignment in the patent text.

#### Compound mismatch

Differences in compound sets mostly reflect extraction coverage. On large patents (*>* 500 PLIs), HARVEST sometimes extracts fewer compounds than BindingDB. On small and medium patents, the opposite holds: HARVEST captures data formats that manual curators skip, including inline activity values in synthesis text, semi-quantitative statements (“IC_50_ *<* 5 *µ*M for all 590 examples”), and non-numeric activity codes (letter grades, symbolic ratings, +/++ scales) used in place of exact measurements. Additional differences arose from HARVEST extraction or structure-normalization errors. Overall, HARVEST yields somewhat fewer PLIs on shared patents (606K vs. 691K), primarily due to incomplete extraction from the largest documents.

#### No overlap

The rare cases (1.6%) where neither compounds nor targets overlap are often caused by BindingDB patent mapping or identifier errors, or by the same target mismatch and compound extraction factors described above.

### C. H-Bench: A Public Benchmark Dataset

A central deliverable of this work is **H-Bench**, an open benchmark of bioactivity data extracted by HARVEST that is not present in BindingDB. BindingDB serves as the primary training source for most modern bioactivity models, either as a direct source [9] or through its integration into the PDB [28]. H-Bench therefore provides a structurally novel held-out resource for rigorous model evaluation on records likely omitted from training sets.

To establish the benchmark’s integrity, we used a graph-based integer linear program (ILP) to maximize structural separation between HARVEST compounds and existing records. This process identified two distinct subpopulations (Fig. 5): the **Valid** subset (*n* = 60, 130, median similarity ≈ 0.47), occupying novel chemical space, and the **Common** subset (*n* = 117, 028, median similarity ≈ 0.68), containing compounds structurally closer to BindingDB entries. We provide a Python script that implements this algorithm, allowing researchers to compare H-Bench against their own training data. This tool automatically identifies and reassigns any similar structures to the **Common** subset, effectively censoring potential data leakage and preventing over-optimistic results during model evaluation.

**FIG. 5:**
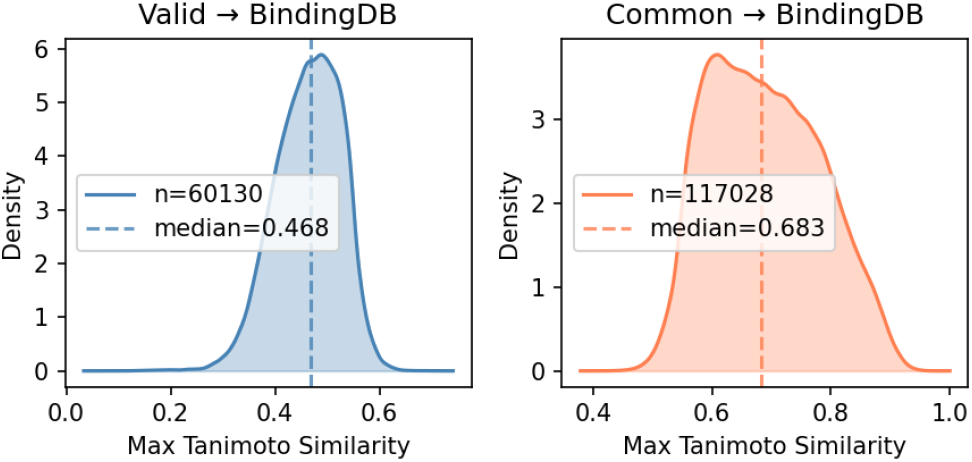
Chemical distance of H-Bench compounds from BindingDB, measured as maximum Tanimoto similarity to the nearest BindingDB compound. (a) *Valid* subset (*n* = 60,130; median = 0.47): compounds with low structural overlap with BindingDB, released as the H-Bench benchmark. (b) *Common* subset (*n* = 117,028; median = 0.68): compounds structurally proximal to BindingDB entries.

H-Bench covers 48 protein targets spanning diverse classes (Supplementary Table S6), including enzymes, membrane receptors, ion channels and transcription factors. Of these, 37 targets overlap with BindingDB and 11 are entirely novel. To ensure reliable evaluation, targets were filtered to maintain an activity balance between 20% and 55% (mean ≈ 33% active, Supplementary Table S6), avoiding the extreme class imbalance common in public screening datasets. These two components, the set of entirely new proteins and the novel structural clusters for overlapping targets, support two primary evaluation scenarios: generalization to novel chemical structures on known targets and generalization to entirely uncharacterized protein targets.

To investigate whether current high-performance models learn transferable binding physics or simply rely on structural similarity to their training data, we evaluated the leading open-source structure-based model Boltz-2 [9] on the H-Bench benchmark. This evaluation uses up to 120 ligands per protein, balanced between active and inactive classes. For the Common subset evaluation, we excluded molecules with Tanimoto similarity above 0.75 to BindingDB to mitigate direct memorization effects.

Of the three Boltz-2 output scores evaluated, `affinity_pred_value` performed best overall. Performance varied systematically across the three benchmark subsets: AUC-ROC increased monotonically from Valid/new targets (novel proteins, novel chemistry; AUC ≈ 0.52) to Valid/known targets (known proteins, novel chemistry; AUC ≈ 0.63) to Common (known proteins, structurally familiar chemistry; AUC ≈ 0.70), directly mirroring proximity to training data (Fig. 6a). The score intended for hit discovery, `affinity_prob_binary`, performed near-randomly across all subsets (AUC ≈ 0.52–0.58), indicating a significant challenge for virtual screening applications where the model must distinguish binders from decoys. The `confidence_score` metric, which reflects structural plausibility of predicted complexes, showed a similar gradient but remained near random on genuinely novel chemistry.

**FIG. 6:**
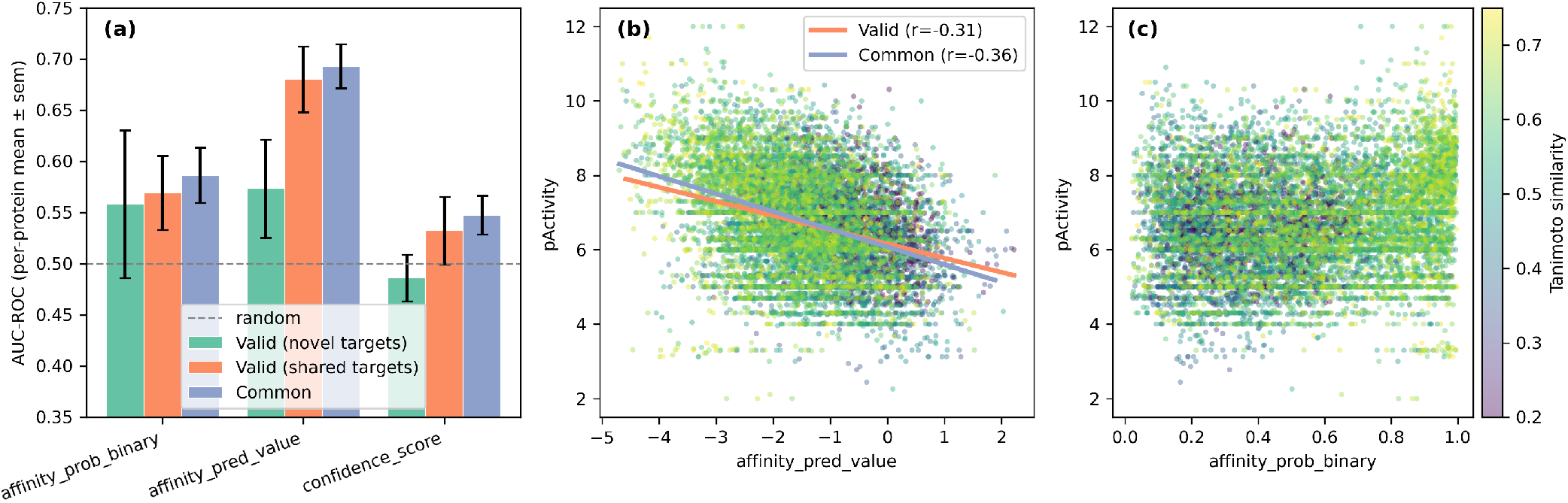
Evaluation of Boltz-2 predicted binding scores against experimental activity on H-Bench. (a) Per-protein AUC-ROC for three Boltz-2 output scores (mean±SEM across proteins), evaluated on three benchmark subsets that differ in both chemical and target novelty: Valid/novel targets (11 proteins exclusive to HARVEST, novel chemistry); Valid/known targets (37 proteins shared with BindingDB, novel chemistry); and Common (37 proteins shared with BindingDB, chemistry structurally proximal to BindingDB). The dashed line marks random performance (AUC = 0.5). Performance increases monotonically with proximity to training data across all three scores, revealing that Boltz-2 predictions reflect structural familiarity more than transferable binding physics. (b) Experimental pActivity (-log_10_[M]) vs. predicted `affinity_pred_value` with linear fits and Pearson r, for Valid/known (orange) and Common (blue) subsets. (c) pActivity vs. `affinity_prob_binary`. Points in (b, c) are colored by Tanimoto similarity to the nearest BindingDB compound.

A key finding is the systematic performance gap between the two subsets: points colored by Tanimoto similarity cluster toward higher predicted scores as similarity increases (Fig. 6b,c). This reveals that model predictions are often biased by proximity to training data rather than underlying binding physics. Scatter analysis confirms only a weak but statistically significant correlation between predicted affinity and experimental activity (Pearson *r* ≈ *−*0.31 for the Valid set).

Overall, Boltz-2 performance on H-Bench is modest but above random on known targets, and degrades on proteins entirely absent from public bioactivity repositories, which is consistent with results reported on the model’s own proprietary benchmarks [9]. This two– dimensional generalization gap — across both chemical and target space — confirms that H-Bench provides a stringent and unbiased evaluation resource. The results suggest that current structure-based models have not yet learned fully transferable binding physics, and that both novel chemical scaffolds and uncharacterized targets remain a fundamental challenge for AI-driven drug discovery.

## III. DISCUSSION

The pharmaceutical industry has invested hundreds of billions of dollars in protein–ligand interaction experiments over the past three decades. Patent law was designed to make this knowledge public, yet the practical inaccessibility of unstructured patent archives has meant that this investment remained effectively private– confined to commercial databases behind expensive subscriptions or simply uncurated. HARVEST transforms this situation: by processing the full USPTO pharmaceutical corpus in under a week at $0.11 per document, it demonstrates that the era of dark bioactivity data is ending. The 3.15 million activity records recovered, including 326, 342 structural clusters and 967 protein targets absent from BindingDB, represent a qualitative expansion of the computable chemical-biological landscape available to the global research community.

This capability distinguishes HARVEST from all existing approaches along two dimensions simultaneously: scale and semantic depth. SureChEMBL provides a high-volume index of chemical structures [21]; however, without quantitative binding context or protein identity resolution it answers “what molecules appear in patents” but not “what do they do and against which target.” BindingDB provides exactly that semantic depth but is fundamentally constrained by the throughput of human expert curation [18]. HARVEST achieves BindingDB-level extraction fidelity – 91% agreement on matched records, with lower unit conversion error rates than manual curators – at 3,500 times the throughput. Recent LLM-based systems like BioMiner [29] and BioChemInsight [30] represent important progress but remain restricted to scientific articles or small-scale proofs-of-concept. HARVEST is the first system to demonstrate this combination of fidelity and scale across a full national patent corpus (Table II).

**TABLE II:** Comparison of automated bioactivity extraction systems. ^*\**^Throughput measured under different conditions: HARVEST uses cloud-based LLM API with 50-document parallelism; BioMiner reports 14s/paper on 8×V100 GPUs. Only HARVEST performs full resolution to UniProt identifiers and standardized units.

|  | HARVEST | BioMiner<br>[29] | BioChemInsight<br>[30] |
| --- | --- | --- | --- |
| Source | Patents | Articles | Patents |
| Documents processed | 164,877 | 11,683 | 181 |
| Documents with bioactivity | 40,902 | 11,683 | N/A |
| Records extracted | 3.15M | 68K | N/A |
| Throughput* | 2 sec/doc | 14 sec/doc | N/A |
| SMILES | Yes | Yes | Yes |
| Protein ID | UniProt | AlphaFold path | N/A |
| Unit normalization | Yes | No | No |
| Public dataset | Partially | Yes | No |

Beyond raw extraction, a significant advantage of HARVEST is its automated data normalization. The system resolves varied and often ambiguous protein descriptors to canonical UniProt identifiers [31] and standardizes diverse activity units into a consistent numeric format. By producing a dataset structured similarly to BindingDB, HARVEST provides a machine-actionable resource that is immediately ready for training machine learning models. This eliminates the massive manual post-processing and data-cleaning efforts typically required when working with “dirty” automated extractions from patent literature.

The cost profile of HARVEST fundamentally reshapes the economics of medicinal chemistry data. Traditional manual curation projects like BindingDB process approximately 1,500 patents over two years [18]. At that rate, processing our full 164,877-patent corpus would require over 55 years of continuous expert labor, whereas HARVEST completed the task in under a week (Table II).

This efficiency removes the financial and structural barriers that have long confined high-volume bioactivity data to proprietary commercial platforms such as Reaxys [32, 33] or GOSTAR [34]. Beyond cost, HARVEST addresses the inherent opacity of commercial databases. While platforms like GOSTAR are often treated as a “black box,” our pipeline provides a transparent, reproducible, and fully auditable lineage from patent XML to final SMILES.

Furthermore, because commercial curation is often driven by market demand for high-traffic therapeutic areas, niche antiviral research or rare-disease data can be deprioritized or excluded. HARVEST captures this “long tail” of chemical space with equal fidelity. Finally, manual curation introduces a lag of years between publication and inclusion. HARVEST can be integrated with weekly USPTO releases, maintaining a near-real-time mirror of the patent landscape—a capability that remains structurally impossible for manual systems regardless of funding.

A striking finding is that despite recovering 967 protein targets entirely absent from BindingDB, these novel targets account for only 2.4% of total extracted interactions. The vast majority of new data deepens coverage of established targets rather than revealing entirely new biological associations. This pattern likely reflects two converging forces: the pharmaceutical industry’s strategic focus on validated targets where biological risk is lower, and the temporal lag between patent filing and scientific publication.

The release of H-Bench addresses a fundamental evaluation problem in AI-driven drug discovery. Because modern models are often trained on the same core public datasets, it is difficult to distinguish genuine generalization from memorization [10, 11]. H-Bench provides a structurally guaranteed held-out resource because its records are derived from patent literature currently absent from BindingDB. Our three-way evaluation of Boltz-2 [9] – across novel targets, known targets with novel chemistry, and known targets with familiar chemistry – reveals that the generalization gap is two-dimensional: models degrade both when chemistry is novel and when targets lack prior bioactivity data. This is a more precise characterization of model limitations than binary train/test splits on BindingDB alone can provide, and it points directly toward what the field needs to improve: training data that covers undercharacterized targets, and evaluation frameworks that separately stress-test chemical and biological generalization. H-Bench provides both.

The success of HARVEST relies heavily on the high quality of structured data provided by the USPTO. Because the USPTO includes chemical structures as embedded ChemDraw files within its XML archives, they can be reliably converted into canonical SMILES. In contrast, processing raw PDF documents would require complex OCR and significantly more intensive processing, which often introduces errors. This highlights the critical importance of maintaining publicly available data in well-organized, machine-readable formats to enable automated discovery.

The 2025 announcement by the CNIPA in China to promote XML for electronic patent submissions [35] opens a direct pathway to extend HARVEST to Chinese pharmaceutical literature, which would add a massive and currently inaccessible reservoir of medicinal chemistry data. More broadly, the value of open, structured data formats for enabling downstream scientific computation cannot be overstated: the difference between a PDF and an XML archive is the difference between dark data and actionable knowledge.

The principles demonstrated by HARVEST extend far beyond bioactivity extraction. Much of humanity’s expert knowledge remains practically inaccessible: technically public in patents, regulatory filings, and clinical records, yet effectively “dark” due to the prohibitive cost of human curation. The multi-agent architecture introduced here, which decomposes complex document understanding into specialized sequential agents grounded by structured resolution pipelines, is directly applicable to any domain facing this barrier. Recent examples include agentic systems for multilingual pharmaceutical asset scouting [36, 37] and automated extraction from clinical and regulatory documents [38, 39]. As LLM capabilities improve and inference costs decline [24], the marginal cost of converting massive document corpora into structured knowledge is approaching zero. The critical question is no longer whether this conversion is feasible, but which datasets are most vital to recover and how to ensure the resulting knowledge remains a public good rather than a proprietary asset.

## IV. LIMITATIONS

HARVEST’s current scope defines a clear roadmap for future development. Four categories of patent data remain to be addressed. First, the system cannot yet process Markush structures. These are highly complex diagrams used to represent large families of molecules simultaneously via variable parts. Automatically “unpacking” these structures into a list of specific, individual compounds is a significant technical challenge that is not yet implemented, although these cases account for only 8% of the data points we excluded (Section IX D). Second, graphical data such as dose–response curves remain inaccessible, preventing the extraction of parameters from assays reported only in figure form. Third, protein target resolution is limited by the scope of the curated Swiss-Prot database [40] and by the inherent ambiguity of multi-subunit complexes like integrins or IL-23. Patents often reference these targets using inconsistent subunit, heterodimer, or domain-level descriptors, making canonical mapping difficult. Finally, LLM safety policies occasionally caused the system to refuse patents targeting high-risk pathogens such as Ebola virus. This has resulted in systematic coverage gaps in certain antiviral research areas, where the model interprets the data extraction as a violation of safety guidelines.

Our cross-validation is grounded in the 5,668 patents shared with BindingDB, representing 14% of the HARVEST corpus. For the remaining 86%, no independent reference currently exists – establishing such a reference, through prospective experimental validation of HARVEST-derived predictions or through community curation efforts, is a priority for future work. Based on the consistency of activity residual distributions between validated and unvalidated subsets, we estimate the error rate for uncharacterized records at level not exceeding 10-15%, comparable to known error rates in manually curated databases [18].

## V. CONCLUSION

We have presented HARVEST, a multi-agent LLM pipeline that converts the dark bioactivity data of pharmaceutical patents into open, computable scientific infrastructure. By decomposing complex document understanding into specialized sequential agents, HARVEST achieves BindingDB-level extraction fidelity at 3,500 times the throughput of manual curation – processing 164,877 patent archives in under a week at $0.11 per document, recovering 326,342 structural clusters and 967 protein targets entirely absent from public repositories. At this cost, comprehensive patent mining is no longer a luxury for well-funded commercial providers; it is accessible to any research group with a compelling scientific question.

The accompanying H-Bench benchmark addresses an equally critical problem: the lack of genuinely heldout evaluation data for protein–ligand interaction models. Our three-way evaluation of Boltz-2 [9] reveals that the generalization gap is two-dimensional – model performance degrades both when chemistry is novel and when protein targets lack prior public bioactivity data. This finding exposes a fundamental limitation of models trained exclusively on manually curated repositories, and establishes H-Bench as a stringent, leakage-free resource for driving the development of more robust, physics-aware models.

Together, HARVEST and H-Bench represent a practical answer to the data bottleneck that currently limits AI-driven drug discovery. The hundreds of billions of dollars in R&D investment embedded in the global patent corpus was always legally public; it was never computationally accessible. As LLM inference costs continue to fall [24], the same approach can be extended to new patent jurisdictions, to regulatory filings, and to any domain where expert knowledge remains trapped in unstructured text. The era of dark data in medicinal chemistry is ending.

## VI. ACKNOWLEDGMENTS

We thank Mikhail Batin, Alexey Strygin, and Vita Stepanova, the organizers of the Agentic AI Against Aging (AAAA) hackathon, for providing the venue that facilitated the inception of this work. We are also deeply grateful to Vladimir Manujlov, an employee of Gero, for his assistance and guidance during the hackathon and for his valuable edits and review of this manuscript. We extend our gratitude to Daniel Kravtsov for providing his technical expertise in modern agentic systems, which offered helpful insights during the design of the HARVEST pipeline. Finally, we acknowledge the use of the Gemini Large Language Model for assistance in the drafting and linguistic refinement of this manuscript.

## VII. CONFLICTS OF INTEREST

K.A., L.M. and P.F. are employees and equity holders of Gero PTE. LTD., a company developing AI-driven drug discovery tools. Gero proposed the patent mining challenge at the AAAA hackathon and may benefit commercially from the methods and datasets described in this work. V.S., A.M., K.C., and N.A.Z. were selected as the winning implementation team by the independent AAAA organizing committee and subsequently contributed to this work under a contract with Gero. All authors declare no other competing financial interests.

## VIII. DATA AVAILABILITY

The H-Bench benchmark and the full HARVEST dataset are available at https://github.com/gero-science/HARVEST under the Creative Commons Attribution 4.0 International (**CC BY 4.0**) license. The ChemDraw binary file reader is available at https://github.com/gero-science/cdx_reader.

This work also utilizes data from the BindingDB opensource database (September 2025 release), which can be accessed at https://www.bindingdb.org. The raw patent application data used for extraction was obtained from the USPTO Patent Application Full Text Data with Embedded TIFF Images (APPDT), available at https://data.uspto.gov/bulkdata/datasets/appdt.

Detailed dataset statistics, protein family distributions, and activity label balance metrics are provided in the Supplementary Information.

## IX. MATERIALS AND METHODS

### A. Patent Data Sources

We evaluated several repositories for large-scale bioactivity extraction, prioritizing structured data formats, API stability, and cost. Alternative sources such as SureChEMBL [21], Google Patents [41], Google Big-Query Patents Public Data [42] and Lens.org [43] were excluded due to limitations in their data formats (see Supplementary Section S2 for details).

**USPTO Bulk Data** [44] was selected as the primary data source, which provides mostly unlimited downloads supported by an API without any additional costs. The structured XML format of the provided documents preserves table hierarchies and chemistry tags, which is essential for accurately linking compounds to their biological targets and activity values. This high-density disclosure is a major advantage of the patent corpus: US patents contain an average of 160 measurements per document compared to only 40 per scientific article [18].

We retrieved weekly archives from the USPTO Application Data (APPDT) using the Bulk Datasets API. To ensure the corpus was rich in extractable PLI data, we applied an initial filter to retain only documents containing either embedded chemical structure attachments (CDX/MOL) or specific bioactivity keywords (e.g., “IC50”, “Ki”, “Kd”, “EC50”) identified via regular expression matching. The keyword filter was optimized for high recall, minimizing false negatives at the cost of increased false positives that are subsequently filtered by downstream agents. This pre-filtering is supported by our empirical observation that patents lacking CDX/MOL attachments rarely contain structured quantitative bioactivity data.

### B. Patent Family Deduplication

A single invention can generate multiple legally distinct application records, such as pre-grant publications, granted patents, continuation or divisional applications, and reissue or reexamination documents. These filings frequently contain identical bioactivity tables and chemical examples, which can lead to redundant data extraction and inflated record counts. To ensure each unique experimental observation is represented only once, we aggregated related filings into *continuity clusters*.

We constructed these clusters by building a directed graph of parent–child relationships retrieved via the USPTO API. To specifically identify content duplication, we restricted the graph edges to priority claims and continuity links: “is a continuation of,” “is a divisional of,” “is a national stage entry of,” and “is a reissue of.” We explicitly excluded “continuation-in-part” (CIP) relationships from this automated deduplication, as CIP filings often introduce new substantive data not present in the parent application.

For each resulting cluster, we selected the most recent document by publication date as the representative record. This strategy ensures the capture of the most complete version of the disclosure, as later filings often include corrected tables or refined IUPAC nomenclature. This deduplication process yielded a 17.7% reduction in the total document volume, resulting in a final corpus of unique inventions for bioactivity extraction.

### C. Multi-Agent Architecture

To extract PLIs from linguistically complex patent documents, we developed a multi-agent architecture consisting of five specialized agents operating in sequence (Fig. 7). We adopted this multi-stage decomposition because LLMs often exhibit performance loss when required to identify heterogeneous data types simultaneously [23]. Our preliminary experiments on 500 patents confirmed this limitation, as a monolithic single-prompt strategy resulted in three systematic failure modes.

**FIG. 7:**
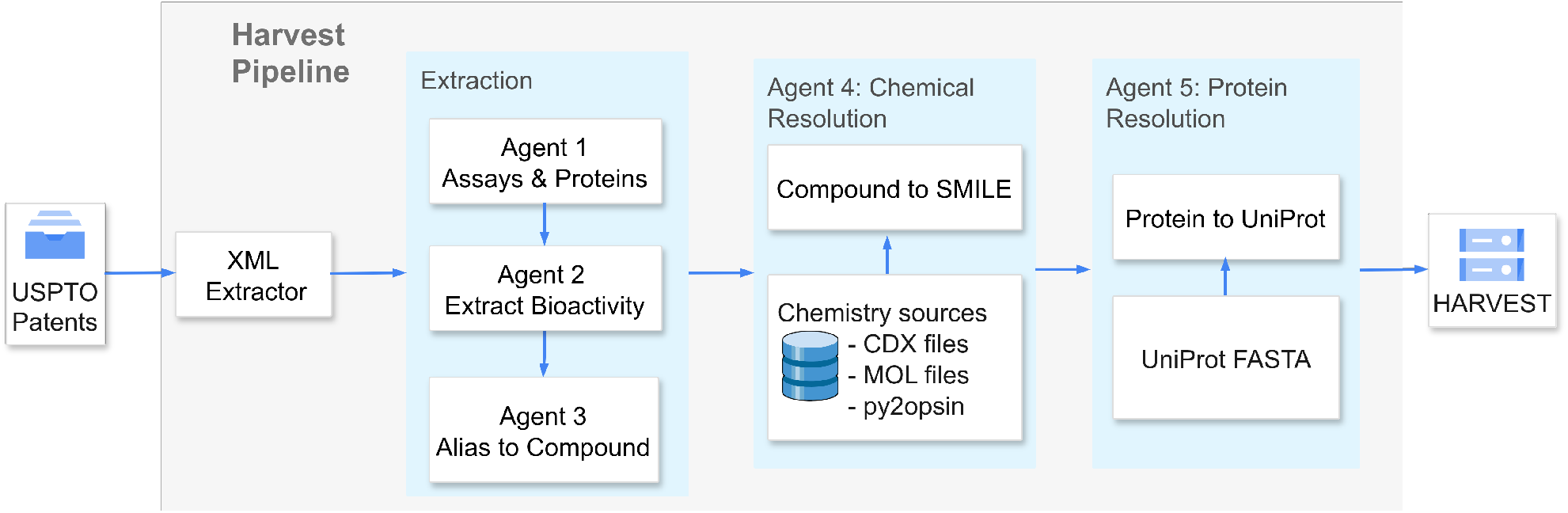
The HARVEST pipeline architecture. USPTO patent documents are parsed by an XML extractor and passed to a three-stage LLM extraction module: Agent 1 identifies biological targets and assay conditions; Agent 2 extracts quantitative bioactivity measurements; and Agent 3 resolves compound aliases to IUPAC names or embedded chemical identifiers. The extracted records are then processed by two resolution agents operating in parallel: Agent 4 converts chemical identities to canonical SMILES via embedded structure files and py2opsin name-to-structure conversion; Agent 5 maps protein names to UniProt identifiers using UniProt FASTA. The final output is the normalized HARVEST dataset of Document–Assay–Result– Compound–Protein records.

First, *attention dilution* caused the model to lose consistent compound–target associations in documents exceeding 200,000 tokens, leading to misattributed activity values. Second, *task interference* substantially increased SMILES hallucination rates when chemical names, biological targets, and numeric values were requested simultaneously. Third, *output truncation* caused premature response termination in patents containing over 500 activity records, losing a substantial fraction of data.

Decomposing the extraction into sequential agents addresses these three failure modes by narrowing the semantic scope of each LLM call. This approach is consistent with recent findings that task decomposition and agent specialization improve performance in frontier models [45–48]. The sequential architecture also improves system traceability and enables targeted prompt optimization for each subtask. Furthermore, this design provides an early termination mechanism: if Agent 1 identifies no biological targets, subsequent steps are skipped to avoid unnecessary computation on irrelevant patents.

The pipeline utilizes three LLM-based agents (Agents 1–3) for semantic extraction, followed by two resolution agents (Agents 4–5) for chemical and protein standardization. All agents operate within an asynchronous processing framework with configurable worker pools, enabling high-throughput parallel processing of the patent corpus.

#### Agents 1–3: Semantic Extraction

**Agent 1 (Target Extraction)** identifies biological targets such as proteins, enzymes, and receptors, along with test organisms and assay conditions. This extracted context is injected into the subsequent Agent 2 prompt to reduce misattribution errors in documents describing multiple targets.

**Agent 2 (Activity Extraction)** focuses on quantitative measurements. It extracts compound aliases (e.g., “Example 1”, “Compound 42”), binding metrics (IC_50_, K_*i*_, K_*d*_, EC_50_), numeric values, and measurement units. By embedding the output of Agent 1, Agent 2 performs target-aware extraction, ensuring that each numeric result is correctly linked to its respective assay and protein.

**Agent 3 (Compound Mapping)** resolves the common patent practice of referencing molecules by internal aliases. It maps these aliases to IUPAC nomenclature or embedded chemical identifiers provided in the text. Isolating this task into a dedicated agent allows handling the complexity: compound identities can be scattered across hundreds of pages, requiring full-document context awareness.

The pipeline utilizes three LLM-based agents to handle the initial semantic extraction from patent text. These agents are built on `google/gemini-2.5-flash`, selected for its 1-million-token context window and efficient prompt caching.

#### Agents 4–5: Resolution

**Agent 4 (Chemical Structure Resolution)** converts resolved chemical identities into standardized SMILES strings. While USPTO patent XML embeds both MOL and CDX (ChemDraw binary) files for all chemical compounds; we use CDX exclusively due to systematic corruption discovered in the distributed MOL representations. This corruption typically occurs during default CDX-to-MOL conversion, where atom type aliases or common substitutions (e.g., “Me”, “Et”, “HN”, or “CN”) are erroneously interpreted as carbon atoms. By parsing the ChemDraw binary files directly, we avoid these substitution errors. The impact of CDX-based resolution on extraction fidelity is shown in Supplementary Table S2. The parser is implemented in Python and is released as an open-source tool (see Data Availability, Section VIII). For patents containing only IUPAC nomenclature without embedded structural files, we use `py2opsin` [49] as a fallback.

**Agent 5 (Protein Target Resolution)** maps extracted protein names to UniProt identifiers [31] and retrieves their associated amino acid sequences. Due to the prevalence of non-standard nomenclature, unofficial aliases, and context-dependent target specification in patent documents, this task requires LLM assistance. We employ `openai/gpt-5-2025-08-07` for this stage, as its superior reasoning capabilities relative to smaller Gemini Flash enable more accurate interpretation of ambiguous biological context. When species is unspecified in the patent text, the agent defaults to *Homo sapiens*, consistent with the therapeutic focus of pharmaceutical patents and the composition of BindingDB, where 83% of binding data derive from human proteins [18].

### D. Dataset Inclusion Criteria and Normalization

HARVEST targets quantitative protein–ligand binding data. The following criteria, enforced through agent prompts and post-processing, define the records included in the final dataset.

#### Binding metrics

Only IC_50_, K_*i*_, K_*d*_, and EC_50_ measurements are retained. These four metrics account for over 80% of records extracted by the agents. Metrics such as percent inhibition are excluded, as extracting meaningful binding constants from single-point or curve-dependent measurements would require additional processing logic not justified by their low prevalence in the corpus. All retained values are normalized to nanomolar (nM).

#### Defined protein targets

Only targets mappable to one or more UniProt identifiers are retained. This includes multi-subunit complexes, which are represented as semicolon-separated accessions. Approximately 87% of extracted records were successfully mapped. Records where the extracted target refers to a cell line or phenotypic endpoint (e.g., “HeLa cytotoxicity” or “A549 cell viability”) rather than a specific protein are excluded.

#### Direct binding assays

Records derived from phenotypic or cell-based readouts, such as cytotoxicity, cell viability, proliferation, are excluded. These reported activity values reflect aggregate cellular responses rather than direct protein–ligand interactions. This filter removed 230,306 records (6.3%).

#### Markush structure exclusion

Patents containing only Markush structures or generic R-group representations are excluded. Enumerating individual compounds from these combinatorial representations requires specialized chemical reasoning not currently implemented in the pipeline. Because most patents additionally report concrete compound examples, this criterion removed only 8.0% of records.

### E. Architectural Design Considerations

Several design decisions shaped the final HARVEST architecture and are described here as they may inform similar extraction systems.

#### Context window and the chunking problem

Our initial architecture chunked patent text to fit within 256K-token context windows. Tables were split into row groups with preserved headers and several surrounding paragraphs for local context. This approach introduced systematic errors: chemical names spanning multiple table cells were truncated at chunk boundaries. More critically, patents frequently define compound aliases (e.g., “Compound 1”) in one section and report bioactivity measurements in distant tables, creating unresolvable cross-references in isolated chunks. The availability of models with 1M+ token context windows resolved these issues entirely, with the additional cost offset by prompt caching, resulting in a 3–5× reduction in per-patent inference cost.

#### Explicit grounding constraints

A persistent failure mode was the generation of plausible but fabricated IUPAC names for compound aliases. Agent 3 was therefore instructed to return chemical identifiers *only* when found verbatim in the patent text, defaulting to TIFF filename references from chemistry tags when no systematic name was available. Combined with downstream validation via py2opsin and CDX file cross-referencing, this constraint reduced false positive chemical identifications in the final dataset.

#### Temperature tuning

Setting the model temperature to 0 was adopted to minimize hallucinated content. Even modest values (e.g., temperature = 0.1) produced noticeably higher rates of fabricated compound names and activity values during our internal validation.

#### Few-shot example leakage

When a patent contained no extractable bioactivity data, the model occasionally output values copied from few-shot prompt examples. We addressed this by engineering the prompts to make “no data found” an explicit valid response and post-processing validation to filter records that matched our prompt examples.

### F. H-Bench Construction

#### Protein selection

We selected 48 proteins spanning diverse target (see Supplementary data). Of these, 37 are “overlap” targets shared with BindingDB (each with *>*30 structural clusters and *>*1,000 BindingDB compounds) and 11 are “novel” targets present only in HARVEST (*>*100 compounds). Diversity across families was ensured by requiring 2–3 representatives per ChEMBL L2 target class and removing redundant proteins via sequence clustering (MMseqs2, 40% identity threshold). An activity-balance filter retained only proteins for which 20–55% of Valid-subset compounds are active.

#### Cluster-based splitting

For each protein, compound Morgan fingerprints (radius 2, 2048 bits) were computed and clustered via complete-linkage hierarchical clustering at a Tanimoto distance threshold of 0.2. Clusters were initially labeled by data source: **A** (HARVEST-only) or **B** (BindingDB-only). A cluster connectivity graph was then constructed using centroid Tanimoto similarity (threshold 0.225, ≤2 hops), and an integer linear program (ILP) identified the minimum set of **A** clusters to relabel as buffer **C**, maximizing structural separation between the Valid (**A**) and BindingDB-only (**B**) subsets.

#### Boltz evaluation

Boltz-2 predictions were generated in affinity mode with 5 recycling steps, MSA server queries, and molecular-weight correction enabled. Up to 120 ligands per protein were sampled, balanced between active and inactive classes and drawn from diverse clusters. For compounds in the Common set, we selected only molecules with Tanimoto similarity below 0.75. Three output scores were evaluated: `affinity_pred_value, affinity_prob_binary`, and `confidence_score`. Per-protein AUC-ROC was computed for proteins with ≥5 compounds and both activity classes present, yielding 5,740 predictions across 48 proteins (Valid) and 4,200 predictions across 37 proteins (Common).

## Supplementary Information

### Appendix S1: Supplementary tables

**TABLE S1:** Distribution of novel protein–ligand interactions (PLIs) and structural clusters contributed by HARVEST, grouped by ChEMBL target classification. For each Level 1 (L1) target class (bold font) containing multiple Level 2 (L2) families, L2 subcategories are listed with indentation. Novel compounds/clusters include all ligands associated with HARVEST-only proteins or ligands not present in BindingDB for shared proteins. Structural clusters were defined using hierarchical complete-linkage clustering of Morgan fingerprints (radius = 2, 2048 bits) with a Tanimoto distance threshold of 0.2, followed by aggregation of adjacent clusters

| Target Class | Proteins | Novel Compounds | Novel Clusters |
| --- | --- | --- | --- |
| **Enzyme** |  |  |  |
| Kinase | 431 | 376,467 | 80,374 |
| Transferase | 253 | 90,049 | 19,130 |
| Protease | 216 | 85,467 | 15,725 |
| Oxidoreductase | 200 | 52,021 | 13,420 |
| Hydrolase | 244 | 50,595 | 10,740 |
| Other | 152 | 47,890 | 12,806 |
| Unspecified | 113 | 24,119 | 5,394 |
| Phosphatase | 61 | 10,174 | 2,922 |
| Lyase | 44 | 1,730 | 545 |
| **Membrane receptor** |  |  |  |
| Family A G protein-coupled receptor | 383 | 219,572 | 43,543 |
| Other | 64 | 50,087 | 10,156 |
| Unspecified | 67 | 16,637 | 5,479 |
| **Ion channel** |  |  |  |
| Voltage-gated ion channel | 97 | 32,784 | 8,439 |
| Ligand-gated ion channel | 53 | 24,449 | 6,751 |
| Other | 18 | 22,111 | 4,723 |
| **Transcription factor** |  |  |  |
| Nuclear receptor | 63 | 52,241 | 8,346 |
| Unspecified | 70 | 19,622 | 7,569 |
| **Unclassified protein** | 306 | 60,665 | 18,073 |
| **Epigenetic regulator** | 108 | 59,705 | 12,714 |
| **Unclassified** | 725 | 31,675 | 13,866 |
| **Transporter** |  |  |  |
| Electrochemical transporter | 95 | 23,640 | 4,975 |
| Other | 33 | 4,983 | 1,867 |
| **Secreted protein** | 78 | 27,374 | 11,017 |
| **Other cytosolic protein** | 54 | 17,556 | 4,128 |
| **Other** | 81 | 15,538 | 3,640 |
| **Total** | **4,077** | **907,692** | **450,625** |

### Appendix S2: Alternative Patent Data Sources

Several alternative patent data sources were evaluated but found unsuitable for corpus-scale bioactivity extraction:

**SureChEMBL** (https://www.surechembl.org) provides free access to patent-extracted chemical structure data through an open API [21]. However, tabular data in the processed output are concatenated without clear delimiters, making it difficult to distinguish decimal separators from column boundaries. The API also exhibits instability under sustained high-throughput querying, returning HTTP 500 errors at the request rates required for corpus-scale extraction.

**Google Patents** (https://patents.google.com) and **Google BigQuery Patents Public Data** were both evaluated but found to be unsuitable: Google Patents implements aggressive rate limiting and IP-based blocking during programmatic access, while BigQuery, though well-structured, proved cost-prohibitive for extracting the complete pharmaceutical patent corpus with full-text descriptions in academic research budgets.

**Lens.org** provides a more cost-effective alternative with institutional subscription options, but patent description fields are provided in plain text rather than structured markup. This formatting limitation often causes tabular data to collapse, where numerical values from separate columns merge into single, unspaced strings. Such degradation prevents the reliable recovery of individual activity values and complicates the resolution of chemical references, both of which are essential for extracting high-fidelity protein–ligand bioactivity relationships.

### Appendix S3: Impact of CDX-based structure resolution

**TABLE S2:** Cross-validation on shared patents without CDX-based chemical structure resolution (using only MOL files and py2opsin). Without the CDX parser, the full-match rate drops from 80.3% to 72.7% for HARVEST records and from 70.5% to 63.6% for BDB records (cf. Table I), while compound mismatches approximately double (from 7.9% to 15.5% and from 18.0% to 25.0%, respectively). These results demonstrate the improvement in extraction fidelity achieved by parsing chemical structures directly from the original ChemDraw binary files.

|  | HARVEST<br>(*n*=603,929)<br>in BDB | BDB<br>(*n*=690,907)<br>in HARVEST |
| --- | --- | --- |
| Full match | 439,113 (72.7%) | 439,113 (63.6%) |
| Target mismatch | 57,079 (9.5%) | 49,151 (7.1%) |
| Compound mismatch | 93,471 (15.5%) | 172,515 (25.0%) |
| No overlap | 14,266 (2.4%) | 30,128 (4.4%) |

### Appendix S4: Cross-validation case studies

**TABLE S3:** Manual review of the largest cross-validation mismatches (a): Target mismatch — the same compound was found in both databases but assigned to different proteins.

| Patent | Target | UniProt ID<br>(HARVEST) | UniProt ID<br>(BindingDB) | Verdict |
| --- | --- | --- | --- | --- |
| US20140275087A1 | GlyT1 | SC6A9_HUMAN | MGAT1_HUMAN | BindingDB incorrectly recognized a protein |
| US20150376212A1 | FLAP | AL5AP_HUMAN | FEN1_HUMAN | BindingDB incorrectly recognized a protein |
| US20240246937A1 | IL4I1 | OXLA_HUMAN | IL1RA_HUMAN | BindingDB incorrectly recognized a protein |
| US20250051309A1 | MK2, p38α | MAPK2_HUMAN,<br>MK14_HUMAN | MK01_HUMAN,<br>4EBP1_HUMAN | BindingDB incorrectly recognized a protein |
| US20170226089A1 | NR2B | NMDZ1_HUMAN;<br>NMDE1_HUMAN;<br>NMDE2_HUMAN;<br>NMDE3_HUMAN;<br>NMDE4_HUMAN;<br>NMD3A_HUMAN;<br>NMD3B_HUMAN | RXRA_HUMAN | HARVEST and BindingDB incorrectly recognized a protein |
| US20150191464A1 | IRAK4 | IRAK4_HUMAN,<br>TLR2_HUMAN | IRAK4_HUMAN | HARVEST incorrectly recognized assay |
| US20140057889A1 | Bcl-XL, Bcl-2 | B2CL1_HUMAN,<br>BCL2_HUMAN | B2CL1_MOUSE,<br>BCL2_MOUSE,<br>BCL2_HUMAN | No information about the organism |
| US20120035149A1 | PKR1, PKR2 | PKR1_HUMAN,<br>PKR2_HUMAN | G3HIE5_CRIGR,<br>G3H407_CRIGR | No information about the organism |
| US20240059703A1 | KRAS G12C,C118A (aa 1-169), His-tagged | RASK_HUMAN | SOS1_HUMAN | HARVEST and BindingDB do not process protein modifications |
| US20170334917A1 | PG-K1 (aa101-181), His-tagged | PLMN_HUMAN | TPA_HUMAN | HARVEST and BindingDB do not process protein modifications |

**TABLE S4:** Manual review of the largest cross-validation mismatches (b): Compound mismatch — the same protein was found in both databases but associated with different compound sets.

| Patent | H | B | Ovlp | Verdict |
| --- | --- | --- | --- | --- |
| US20230255941A1 | 44 | 953 | 33 | HARVEST incomplete extraction |
| US20200095247A1 | 57 | 804 | 56 | HARVEST incomplete extraction |
| US20130172317A1 | 68 | 2136 | 52 | HARVEST incomplete extraction |
| US20220175775A1 | 79 | 1080 | 79 | HARVEST incomplete extraction |
| US20140094448A1 | 205 | 1116 | 179 | HARVEST incomplete extraction |
| US20240208941A1 | 89 | 332 | 89 | HARVEST incomplete extraction |
| US20190322661A1 | 32 | 462 | 30 | HARVEST incomplete extraction |
| US20240294506A1 | 2 | 499 | 2 | HARVEST incomplete extraction |
| US20250129020A1 | 201 | 454 | 197 | HARVEST incomplete extraction |
| US20240294524A1 | 84 | 783 | 68 | HARVEST incomplete extraction<br>(activity table not present in the XML package) |
| US20240254137A1 | 37 | 892 | 4 | Mixed case: HARVEST incomplete extraction<br>and non-matching source documents |
| US20190241588A1 | 754 | 766 | 59 | HARVEST chemical structure error |
| US20250017938A1 | 781 | 1 | 1 | BindingDB incomplete curation |
| US20160304519A1 | 515 | 48 | 48 | BindingDB incomplete curation |
| US20220041601A1 | 284 | 8 | 4 | BindingDB incomplete curation |
| US20120058988A1 | 562 | 94 | 92 | BindingDB incomplete curation |
| US20180118724A1 | 332 | 259 | 146 | HARVEST extraction error |
| US20130079303A1 | 322 | 318 | 115 | HARVEST chemical structure error |
| US20180273529A1 | 443 | 298 | 40 | HARVEST chemical structure error |
| US20230286970A1 | 393 | 405 | 392 | Different inclusion threshold for inactive measurements |

**TABLE S5:** Manual review of the largest cross-validation mismatches (c): No overlap — neither compounds nor targets matched between databases for the same patent.

| Patent | H | B | Verdict |
| --- | --- | --- | --- |
| US20180118720A1 | 105 | 42 | BindingDB patent mapping error |
| US20210009607A1 | 138 | 25 | BindingDB patent mapping error |
| US20240343719A1 | 117 | 20 | BindingDB patent mapping error |
| US20200157106A1 | 5 | 800 | BindingDB patent mapping error |
| US20160115131A1 | 97 | 516 | BindingDB patent mapping error |
| US20210053946A1 | 8 | 612 | BindingDB accession or identifier error |
| US20190343826A1 | 37 | 608 | BindingDB accession or identifier error |
| US20210188777A1 | 163 | 7 | BindingDB incomplete curation |
| US20240209001A1 | 103 | 10 | BindingDB incomplete curation |
| US20220315606A1 | 429 | 3 | BindingDB incomplete curation |
| US20240400523A1 | 1 | 619 | HARVEST incomplete extraction |
| US20250064789A1 | 16 | 540 | Mixed case: identifier mismatch and<br>HARVEST incomplete extraction |
| US20160060224A1 | 63 | 505 | Technical identifier limitation |
| US20230365594A1 | 214 | 8 | Different assay focus across sources |

## Notes

https://github.com/gero-science/HARVEST

https://github.com/gero-science/cdx_reader

### Competing Interest Statement

The authors have declared no competing interest.

### Summary of Updates

Updated abstract, Figure 1, 2, 5 and 6 revised, Figure 7 deleted. Supplementary table S1 updated.

## References

[1] M. Bregonje, Patents: a unique source for scientific technical information in chemistry related industry?, World Patent Inf 27, 309 (2005).

[2] M. E. Valentinuzzi, Patents and scientific papers: Quite different concepts: The reward is found in giving, not in keeping [retrospectroscope], IEEE Pulse 8, 49 (2017).

[3] C. Southan, P. Varkonyi, K. Boppana, S. A. R. P. Jagarlapudi, and S. Muresan, Tracking 20 years of compound-to-target output from literature and patents, PLOS ONE 8, e77142 (2013).

[4] J. L. Watson, D. Juergens, N. R. Bennett, B. L. Trippe, J. Yim, H. E. Eisenach, W. Ahern, A. J. Borst, R. J. Ragotte, L. F. Milles, B. I. M. Wicky, N. Hanikel, S. J. Pellock, A. Courbet, W. Sheffler, J. Wang, P. Venkatesh, Sappington, S. Vázquez Torres, A. Lauko, V. De Bortoli, E. Mathieu, S. Ovchinnikov, R. Barzilay, T. S. Jaakkola, F. DiMaio, M. Baek, and D. Baker, De novo design of protein structure and function with RFdiffusion, Nature 620, 1089 (2023).

[5] N. R. Bennett, J. L. Watson, R. J. Ragotte, A. J. Borst, D. L. See, C. Weidle, R. Biswas, Y. Yu, E. L. Shrock, R. Ault, P. J. Y. Leung, B. Huang, I. Goreshnik, J. Tam, K. D. Carr, B. Singer, C. Criswell, B. I. M. Wicky, D. Vafeados, M. Garcia Sanchez, H. M. Kim, S. Vázquez Torres, S. Chan, S. M. Sun, T. T. Spear, Y. Sun, K. O’Reilly, J. M. Maris, N. G. Sgourakis, R. A. Melnyk, C. C. Liu, and D. Baker, Atomically accurate de novo design of antibodies with RFdiffusion, Nature 649, 183 (2026).

[6] F. J. Ferreira and A. S. Carneiro, Ai-driven drug discovery: a comprehensive review, ACS omega 10, 23889 (2025).

[7] J. Jumper, R. Evans, A. Pritzel, T. Green, M. Figurnov, O. Ronneberger, K. Tunyasuvunakool, R. Bates, A. Žídek, A. Potapenko, A. Bridgland, C. Meyer, S. A. A. Kohl, A. J. Ballard, A. Cowie, B. Romera-Paredes, S. Nikolov, R. Jain, J. Adler, T. Back, S. Petersen, D. Reiman, E. Clancy, M. Zielinski, M. Steinegger, M. Pacholska, T. Berghammer, S. Bodenstein, D. Silver, O. Vinyals, A. W. Senior, K. Kavukcuoglu, P. Kohli, and D. Hassabis, Highly accurate protein structure prediction with AlphaFold, Nature 596, 583 (2021).

[8] J. Abramson, J. Adler, J. Dunger, R. Evans, T. Green, A. Pritzel, O. Ronneberger, L. Willmore, A. J. Ballard, J. Bambrick, S. W. Bodenstein, D. A. Evans, C.-C. Hung, M. O’Neill, D. Reiman, K. Tunyasuvunakool, Z. Wu, A. Žemgulytė, E. Arvaniti, C. Beattie, O. Bertolli, A. Bridgland, A. Cherepanov, M. Congreve, A. I. Cowen-Rivers, A. Cowie, M. Figurnov, F. B. Fuchs, H. Gladman, R. Jain, Y. A. Khan, C. M. R. Low, K. Perlin, A. Potapenko, P. Savy, S. Singh, A. Stecula, A. Thillaisundaram, C. Tong, S. Yakneen, E. D. Zhong, M. Zielinski, A. Žídek, V. Bapst, P. Kohli, M. Jaderberg, D. Hassabis, and J. M. Jumper, Accurate structure prediction of biomolecular interactions with AlphaFold 3, Nature 630, 493 (2024).

[9] S. Passaro, G. Corso, J. Wohlwend, M. Reveiz, S. Thaler, V. R. Somnath, N. Getz, T. Portnoi, J. Roy, H. Stark, D. Kwabi-Addo, D. Beaini, T. Jaakkola, and R. Barzilay, Boltz-2: Towards accurate and efficient binding affinity prediction, bioRxiv 10.1101/2025.06.14.659707 (2025), preprint.

[10] P. Tossou, C. Wognum, M. Craig, H. Mary, and E. Noutahi, Real-world molecular out-of-distribution: Specification and investigation, Journal of Chemical Information and Modeling 64, 697 (2024).

[11] H. Fooladi, T. N. L. Vu, M. Mathea, and J. Kirchmair, Evaluating machine learning models for molecular property prediction: Performance and robustness on out-of-distribution data, Journal of Chemical Information and Modeling 65, 9871 (2025), pMID: 40947919.

[12] D. van Tilborg, L. Rossen, and F. Grisoni, Molecular deep learning at the edge of chemical space, ChemRxiv 10.26434/chemrxiv-2025-qj4k3-v3 (2025), preprint.

[13] N. Pallikkavaliyaveetil and S. Chandrasekaran, Small data, big challenges: Machine- and deep-learning strategies for data-limited drug discovery, Advanced Drug Delivery Reviews 10.1016/j.addr.2025.115762 (2025).

[14] K. Debnath, P. Rana, and P. Ghosh, A survey on deep learning for drug-target binding prediction: models, benchmarks, evaluation, and case studies, Briefings in Bioinformatics 26, bbaf491 (2025).

[15] A. Gangwal, A. Ansari, I. Ahmad, A. K. Azad, and W. M. A. W. Sulaiman, Current strategies to address data scarcity in artificial intelligence-based drug discovery: A comprehensive review, Computers in Biology and Medicine 179, 108734 (2024).

[16] J. Dagdelen, A. Dunn, S. Shen, et al., Structured information extraction from scientific text with large language models, Nat Commun 15, 1418 (2024).

[17] J. Michels et al., Natural language processing methods for the study of protein–ligand interactions, J Chem Inf Model 65, 2191 (2025).

[18] T. Liu, L. Hwang, S. K. Burley, C. I. Nitsche, C. Southan, W. P. Walters, and M. K. Gilson, Bindingdb in 2024: a fair knowledgebase of protein-small molecule binding data, Nucleic acids research 53, D1633 (2025).

[19] D. M. Jessop, S. E. Adams, and P. Murray-Rust, Mining chemical information from open patents, J Cheminform 3, 40 (2011).

[20] M. Krallinger, F. Leitner, O. Rabal, M. Vazquez, J. Oyarzabal, and A. Valencia, Chemdner: the drugs and chemical names extraction challenge, J Cheminform 7, S1 (2015).

[21] G. Papadatos, M. Davies, N. Dedman, et al., Surechembl: a large-scale, chemically annotated patent document database, Nucleic Acids Res 44, D1220 (2016).

[22] Y. Gadiya, S. Shetty, M. Hofmann-Apitius, P. Gribbon, and A. Zaliani, Exploring surechembl from a drug discovery perspective, Scientific data 11, 507 (2024).

[23] S. Qiao, R. Fang, Z. Qiu, X. Wang, N. Zhang, Y. Jiang, P. Xie, F. Huang, and H. Chen, Benchmarking agentic workflow generation, arXiv preprint 2410.07869 (2024).

[24] G. Appenzeller, Llmflation: Llm inference cost going down fast, https://a16z.com/llmflation-llm-inference-cost/ (2024), discusses Moore’s-law-like declines in LLM inference cost at fixed quality.

[25] A. L. Hopkins and C. R. Groom, The druggable genome, Nature reviews Drug discovery 1, 727 (2002).

[26] D. Stumpfe, H. Hu, and J. Bajorath, Evolving concept of activity cliffs, ACS Omega 4, 14360 (2019).

[27] S. R. Heller, A. McNaught, I. Pletnev, S. Stein, and D. Tchekhovskoi, Inchi, the iupac international chemical identifier, J Cheminform 7, 23 (2015).

[28] H. M. Berman, J. Westbrook, Z. Feng, G. Gilliland, T. N. Bhat, H. Weissig, I. N. Shindyalov, and P. E. Bourne, The protein data bank, Nucleic Acids Research 28, 235 (2000).

[29] J. Yan, J. Zhu, Y. Yang, Q. Liu, K. Zhang, Z. Zhang, X. Liu, B. Zhang, K. Gao, J. Xiao, et al., Biominer: A multi-modal system for automated mining of protein-ligand bioactivity data from literature, bioRxiv, 2025 (2025).

[30] Z. Wang, F. Fu, W. Zhang, L. Yan, Y. Meng, J. Wu, H. Wu, G. Xu, and S. Chen, Biocheminsight: An open-source toolkit for automated identification and recognition of optical chemical structures and activity data in scientific publications, arXiv preprint 2504.10525 (2025).

[31] The UniProt Consortium, Uniprot: the universal protein knowledgebase in 2023, Nucleic Acids Res 51, D523 (2023).

[32] Elsevier, Reaxys, https://www.elsevier.com/product s/reaxys (2026), accessed: 22 February 2026.

[33] Elsevier, Overview of reaxys data, https://www.elsevier.support/dataasaservice/answer/overview-of-reaxys-data (2026), accessed: 22 February 2026.

[34] Excelra, Gostar™ small molecules, https://www.excelra.com/databases/gostar (2026), accessed: 22 February 2026.

[35] China National Intellectual Property Administration, Promoting the use of extensible markup language (xml) format for submitting electronic patent application documents, https://www.cnipa.gov.cn/art/2025/5/26/art_75_199841.html (2025), accessed: 2026-01-25.

[36] A. Vinogradova, V. Vinogradov, L. Greenwood, I. Yasny, D. Kobyzev, S. Kasbekar, K. Nguyen, D. Radkevich, R. Doronin, and A. Doronichev, Hunt globally: Wide search AI agents for drug asset scouting in investing, business development, and competitive intelligence, arXiv preprint 2602.15019 (2026), preprint.

[37] A. Vinogradova, V. Vinogradov, D. Radkevich, I. Yasny, D. Kobyzev, I. Izmailov, K. Yanchanka, R. Doronin, and A. Doronichev, LLM-based agents for competitive landscape mapping in drug asset due diligence, arXiv preprint 2508.16571 (2025), preprint.

[38] L. Liu, V. Blake, M. Barman, B. Gallego, T. Churches, G. Kennedy, S.-Y. Ooi, G. P. Delaney, and L. Jorm, Using natural language processing to extract information from clinical text in electronic medical records for populating clinical registries: a systematic review, Journal of the American Medical Informatics Association 33, 484 (2026).

[39] Z. Yang, H. Yuan, R. Sayeed, A. L. M. Tan, E. Cai, M. Moro, X. Li, H. Ying, N. Brown, G. Weber, et al., CLINES: Clinical LLM-based information extraction and structuring agent, medRxiv, 2025 (2025), preprint.

[40] E. Boutet, D. Lieberherr, M. Tognolli, M. Schneider, and A. Bairoch, Uniprotkb/swiss-prot, Methods in molecular biology (Clifton, N.J.) 406, 89 (2007).

[41] Google, Google patents, https://patents.google.com/ (2026), accessed: 22 February 2026.

[42] Google, Google patents public datasets: connecting public, paid, and private patent data, https://cloud.goog le.com/blog/topics/public-datasets/google-patents-public-datasets-connecting-public-paid-and -private-patent-data/ (2017), accessed: 22 February 2026.

[43] The Lens, Lens.org, https://www.lens.org/lens/user/subscriptions (2026), accessed: 22 February 2026.

[44] USPTO, Uspto bulk data directory, https://data.uspto.gov/bulkdata/datasets (2026), accessed: 22 February 2026.

[45] T. Zhou, J. Xu, G. Liu, J. Liu, H. Wang, and E. Wu, An approach for systematic decomposition of complex LLM tasks, arXiv preprint 2510.07772 (2026), introduces ACONIC framework showing decomposition improves agent performance on complex tasks.

[46] S. Montazeri, Y. Feng, and K. Sha, PublicAgent: Multiagent design principles from an LLM-based open data analysis framework, arXiv preprint 2511.03023 (2025), shows specialization provides value independent of model strength with 97.5% agent win rates.

[47] Y. Yang, Q. Peng, J. Wang, Y. Wen, and W. Zhang, LLM-based multi-agent systems: Techniques and business perspectives, arXiv preprint 2411.14033 (2024), discusses dynamic task decomposition and organic specialization advantages over single-agent systems.

[48] U.-e. Nisa et al., Agentic ai: The age of reasoning—a review, Journal of Artificial Intelligence (2025), a survey of agentic AI systems and their reasoning capabilities.

[49] D. M. Lowe, P. T. Corbett, P. Murray-Rust, and R. C. Glen, Chemical name to structure: Opsin, an open source solution, J Chem Inf Model 51, 739 (2011).

